# Decoding continuous goal-directed movement from human brain-wide intracranial recordings

**DOI:** 10.1101/2025.02.05.636287

**Authors:** Maarten C. Ottenhoff, Maxime Verwoert, Sophocles Goulis, Simon Tousseyn, Johannes P. van Dijk, Maryam M. Shanechi, Omid G. Sani, Pieter Kubben, Christian Herff

## Abstract

Reaching out your hand is an effortless yet complex behavior that is indispensable in daily life. Restoring arm functionality is therefore rated a top priority by people with tetraplegia. Recently, neural correlates of movement have been observed and decoded beyond the motor cortex, but the degree and granularity of movement representation is not fully understood. Here, we explore the neural content of brain-wide movement-related neural activity by decoding neural correlates into 12 different kinematics of goal-directed reaching behavior. Eighteen participants implanted with stereotactic electroencephalography electrodes performed a gamified 3D goal-directed movement task. We demonstrate that continuous movement kinematics can be decoded from distributed recordings using low, mid and high frequency information in all participants using preferential subspace identification (PSID). The neural correlates of movement were distributed throughout the brain, including deeper structures such as the basal ganglia and insula. Moreover, we show that hand position could only be decoded using a goal-directed reference frame, indicating that widespread low-frequency activity is involved in higher-order processing of movements. Our results strengthen the evidence that widespread motor-related dynamics exist across numerous brain regions and can be used to continuously decode movement. The results may provide new opportunities for motor brain-computer interfaces for individuals with a compromised motor cortex, e.g. after stroke, or for control signals in adaptive closed-loop systems.

## 1. Introduction

Reaching your arm out to grab an object is an effortless, yet complex goal-directed behavior. The functional utility of goal-directed movements is indispensable, and consequently rated top-priority to restore by those who lost the ability to reach [1– 3]. The focus to restore movement using brain-computer interfaces (BCIs) has been on the motor cortex [4, 5], but a new perspective is arising that the whole brain is involved during the generation and control of reaching movement [6]. Decoding these global neural patterns may provide new opportunities to enhance current motor BCIs or for the next generation of brain-computer interfaces targeting people with compromised motor cortices, for example for people suffering from a stroke. To this end, reports in animals and humans observe movement-related neural correlates throughout the animal and human brain [6–10] supporting the view that nearly the whole brain is engaged to generate goal-directed movements. Moreover, these neural correlates have been utilized to predict multiple individual movement parameters, including different gestures, directions, reaches and grasps [11–20].

However, it is unknown to what degree continuous goal-directed reaching movement trajectories can be decoded from brain-wide recordings. By leveraging the unique distributed properties of stereotactic encephalographic recordings (sEEG) [21], studies have demonstrated successful decoding one or two dimensional trajectories of individual movement parameters from cortical and subcortical areas, including continuous force decoding [22, 23] and hand flexion [17, 24]. When broadening the scope to discrete movement decoding using sEEG, studies have reported significant decoding of different grasp types [14], directions in a center out task [19] or force levels [25]. The neural inputs for these decoders span a wide range of frequencies, including low frequencies such as the local motor potential, delta power or theta power [14, 16, 19, 26], mid frequencies including alpha and beta powers [11, 14, 17–19, 27] and high frequency broadband high-gamma power [11, 14, 15, 17, 18, 27, 28], suggesting that multiple movement-related processes at different scales are at play and can be decoded into a movement output [29, 30]. Specifically, low frequency activity and high-gamma power are reported to yields substantial correlates with multiple movement parameters [31]. Particularly the phase of low frequency activity may be related to ongoing goal-directed behavior. Decoding a low-pass filtered (< 3*Hz*) signal led to the highest decoding performance in a 2D center-out task as opposed to using the low frequency power [28]. Using this low frequency signal also shares similarities with the local motor potential reported in the motor cortex [32].

The decoding of several individual movements calls for a comprehensive overview of goal-directed movements. Thus, in this work we aimed to decode 12 different kinematics from distributed sEEG recordings in a continuous 3-dimensional goal-directed reaching task. We recorded neural data from 18 epilepsy patients, covering 119 unique brain areas with 1903 contacts. We included low, mid and high frequency information and used preferential subspace identification (PSID) [33] to decode different behavior kinematics from the neural signal. The decoder was able to reconstruct non-directional hand movement speed significantly above chance for nearly all participants and all frequency information, regardless of electrode placement. Directional hand movement speed could be decoded using low frequency phase information. Our results demonstrate that the neural correlates of the kinematics used to describe goal-directed reaching movements are distributed throughout the brain. Finally, we demonstrate that the position of the hand can only be decoded by changing the reference frame from an allocentric reference frame to a goal-directed reference frame.

## 2. Methods

### Participants

We included 18 epilepsy patients implanted with sEEG electrodes. All participants were under treatment for medication-resistant epilepsy and undergoing an assessment period in preparation for resection surgery. Each participant was implanted with 5 to 14 electrode shafts, resulting in a total of 156 electrode shafts with 1903 contacts over all participants. The implantation locations were determined solely by their clinical need and were not influenced in any way by this study.

### Ethical Approval

The experimental protocol was approved by the institutional review board of Maastricht University and Epilepsy Center Kempenhaeghe (METC20180451). All experiments were in accordance with local guidelines and regulation and were under supervision of experienced healthcare staff. All participants joined the study voluntarily and provided written informed consent.

### Experiment

We developed a continuous 3D movement task where the participant was instructed to capture targets on screen by moving the cursor to the target position (Figure 1f, i). The cursor was controlled by moving their preferred hand above a motion tracker. The tracker fits a hand model on the images of two infrared cameras inside the motion tracker. The coordinates of the hand palm, provided by the fitted hand model, were mapped to the coordinates of the game, enabling the participant to move the cursor. The x and y axis were mapped to left-right and up-down movements, respectively. To represent the third dimension on a 2D screen, we mapped the forward-backward movements to the size of the cursor: moving forward reduced the size of the cursor, whereas moving backward (towards the body) increased the size of the cursor. To capture the target, the participant was instructed to move the cursor to the target and hold it there for one second. The cursor turned green when the location and size were matched correctly with the target. The trial was paused when the hand of the participant was out of view.

**Figure 1.**
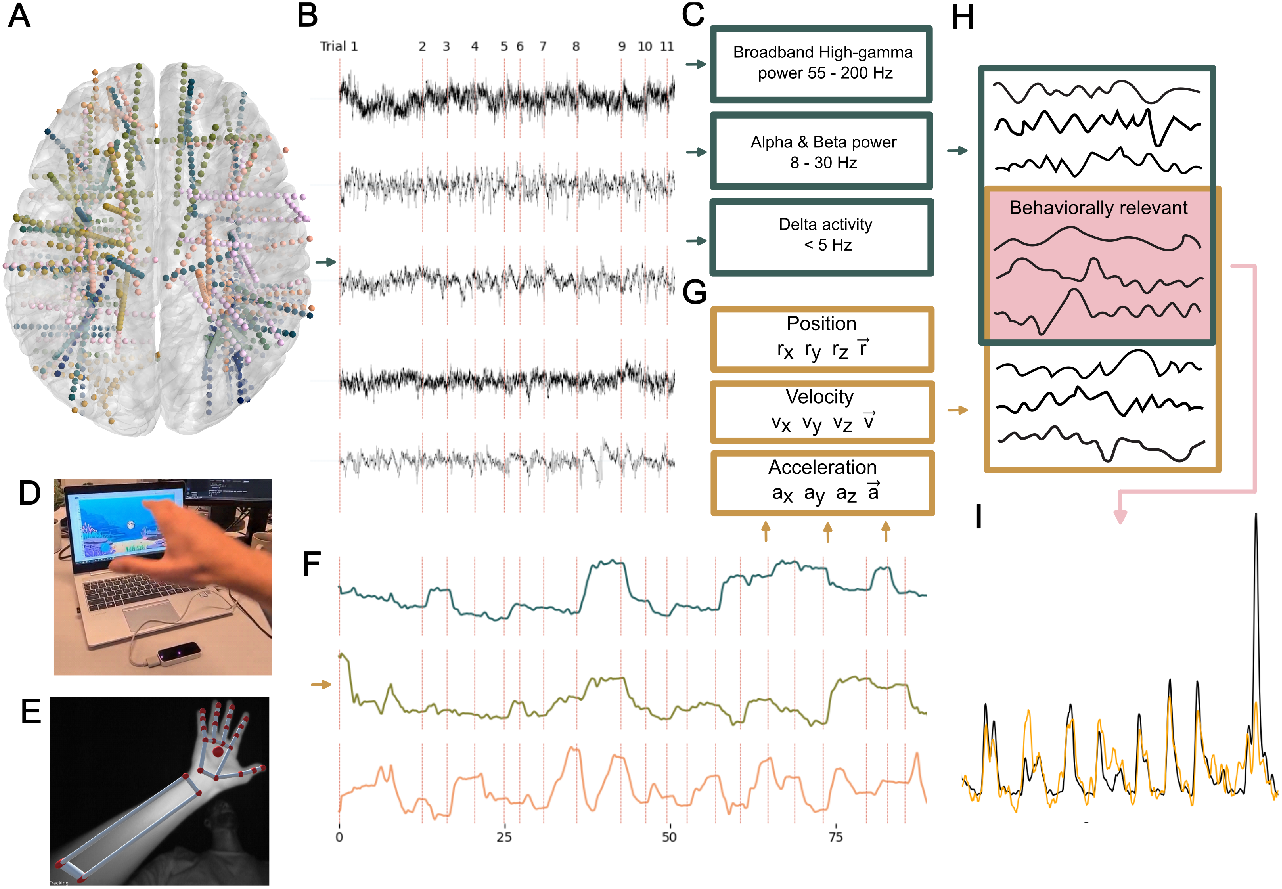
A) Neural Data: Electrode configurations for all participants warped onto an average brain. Each color represents the electrodes of one participant. **B)** Raw EEG data labeled per trial. **C)** Frequency information extracted from the neural data. Delta activity was acquired using a low-pass filter with a cutoff frequency of 5 Hz. Alpha-beta and broadband high-gamma power was extracted by a band-pass filter of 8-30 and 80-200 Hz, respectively, followed by a Hilbert transform. **D) Experiment:** The participants played a game where goals had to be acquired by moving the cursor to the goal and holding it there for 1 second. The cursor was represented as a pufferfish and the goal as a bubble that needed to be popped. The cursor was controlled by a LeapMotion controller (black device in front of PC). Depth (forward-backward movements) was represented in the game by an increase or decrease of the cursor size. For example, to capture a small target, the participant had to move the hand forward to decrease the cursor size such that it matched the size of the target. **E)** To control the cursor, the LeapMotion fits a hand model to images of the hand. The hand palm (large red sphere, middle of hand) was used to control the cursor. **F)** The LeapMotion generated continuous 3D coordinates of the hand palm, where the reference frame was centered at the LeapMotion. **G)** Twelve hand kinematics were extracted from the 3D hand trajectories: the x, y, z coordinates of position *r*, velocity *v* and acceleration a, as well as the non-directional distance (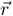, speed 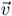 and acceleration 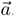. **H) Decoding:** The PSID algorithm was used to extract the behaviorally relevant states from the neural data and hand-movement kinematics. The decoder was trained using five-fold cross validation, during which the 30 most correlated features were selected using four-fold inner cross validation. **I)** The decoder predicted all twelve kinematics, which were evaluated by the correlation between the real (black) and predicted (orange) kinematics. The trajectories shown here represent the predicted speed 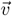 for one of the participants.

When paused, a red target was shown in the middle of the screen. The participant could resume the trial by ‘capturing’ the red target, by moving the cursor to the target. This allowed the participant to take a break during the game, if needed. Targets were presented sequentially and aimed to have a uniform distribution of distances between targets. However, the coordinates were constrained by the box where the motion tracker was able track the hand, and long distances between targets did not always fit. Hence, the distance distribution favored shorter distances over larger distances between targets. To make the task more engaging for our participants, we changed the visual elements to represent an underwater world, where a pufferfish (the cursor) had to pop the bubbles (targets). We captured as much data per participant as recording time and participant ability allowed. For quick and skillful participants, we were able to do multiple runs of 50 targets.

### Recording Setup

Participants were implanted with platinum-iridium sEEG electrodes (Microdeep intracerebral electrodes; Dixi Medical, Beçanson, France), where each electrode shaft contained 5 to 18 contacts. The contacts were 2mm long, 0.8mm in diameter and had an intercontact distance of 1.5mm. The neural activity was recorded using two stacked Micromed SD LTM amplifiers (Micromed, S.p.A., Treviso, Italy). Contacts were referenced to a white matter electrode that did not show epileptic activity, determined visually by the epileptologist. Throughout this work, we use the term contact to refer to the physical recording location on the electrode shaft, and use the term channel to refer to the digitized data from a single contact. We recorded hand movements by using the UltraLEAP motion tracker (Ultraleap Limited, Bristol, England). The tracker records data varying sample rate (effective frame rate = 65 ± 14 Hz). Neural, stimuli and hand movement data streams were synchronized using LabStreamingLayer [34] and recorded using T-Rex [35].

### Imaging, Anatomical labeling & Visualization

The anatomical locations of each contact were determined by co-registering a preimplantation anatomical T1-weighted MRI scan with a post-implantation CT scan. The MRI scan was automatically parcellated in Freesurfer(https://surfer.nmr.mgh.harvard.edu/) according to the Destrieux atlas [36], while contact locations and their corresponding anatomical labels were extracted via the img pipe toolbox [37]. To visualize electrode configurations across participants in a single brain, we warped the native brain and electrode coordinates to the CVS average-35 atlas in MNI512 space. The anatomical labels were always determined in native space. Contacts localized outside of the brain, such as those not inserted, located in sulci or in previously resected brain areas were labeled as unknown. The color maps used throughout this work use the perceptually uniform and color-vision deficiency friendly color maps by Crameri F. (2018) [38, 39].

### Data preparation

To prepare the data for our decoder, in short, we first applied a common electrode re-reference (all contacts on one electrode shaft). Then we extracted low, mid or high frequency information (Figure 1 & 2). To evaluate the spatial correlations, we replaced the common electrode re-reference with a Laplacian re-reference in order to attenuate the effects of passive volume conduction (Figure 3). We next explain the details of these steps.

**Figure 2.**
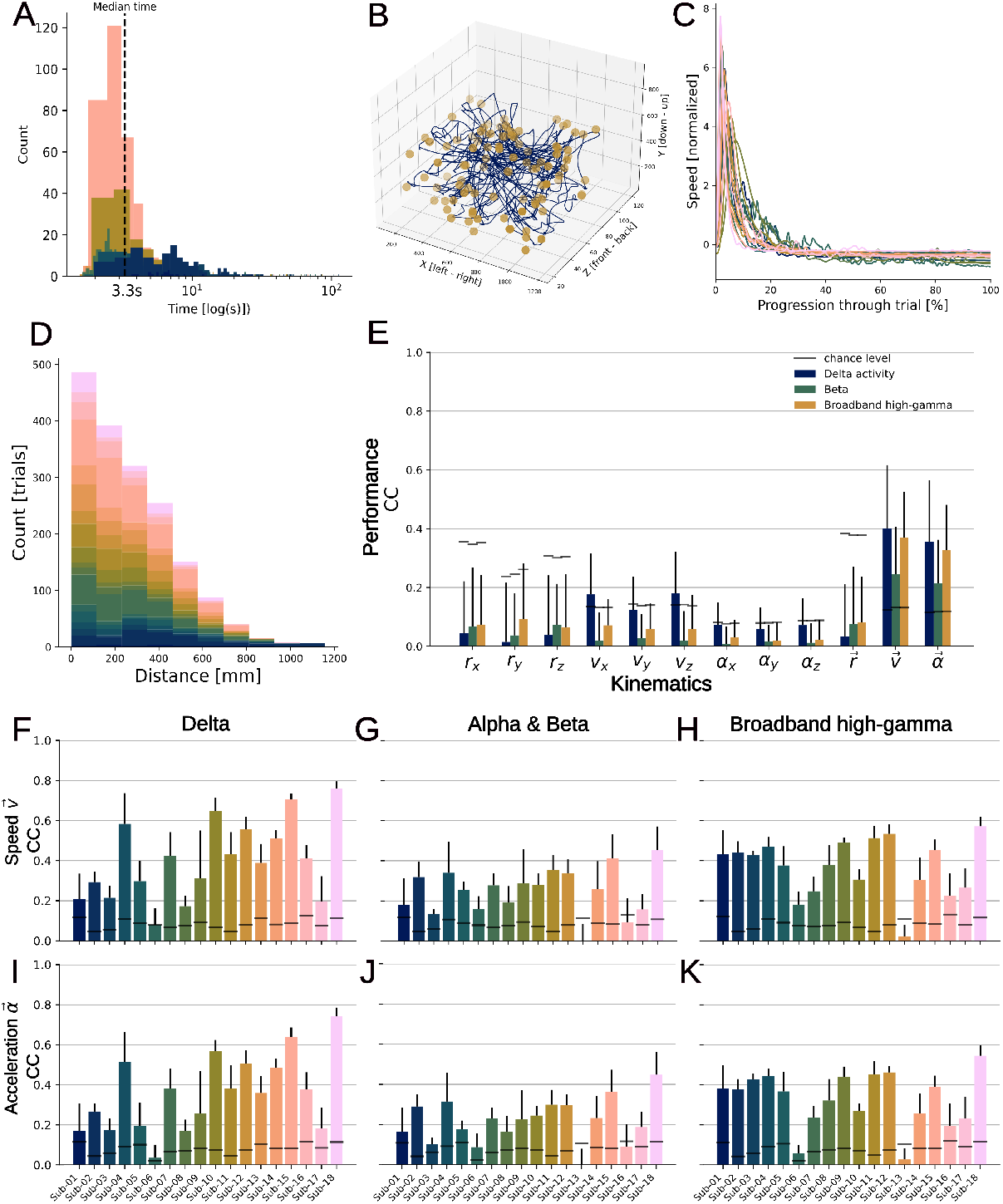
**A)** Time to completion per trial, colored by participant. **B)** Movement trajectory (blue) for one example participant that successfully captured 100 targets (gold circles). **C)** Normalized average speed curve per participant. **D)** Distribution of distances between sequential targets. **E)** Mean decoding performance over participants per kinematic and frequency information. Our decoder was able to reconstruct non-directional speed 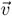 and acceleration 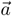 significantly above chance using low, mid and high frequency information. *v*_*x*_, *v*_*y*_ and *v*_*z*_ could be decoded significantly above chance as well. Chance level was determined by n = 1000 circular shift permutations. **F** Reconstruction correlation for velocity 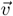 per participant for delta activity. **G)** Same as **F**, but for alpha-beta power. **H)** Same as **F**, but for broadband high-gamma power. **I)** Reconstruction correlation for acceleration 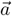 per participant for delta activity. **J** Same as **I**, but for alpha-beta power. **K)** Same as **I**, but for broadband high-gamma power.

**Figure 3.**
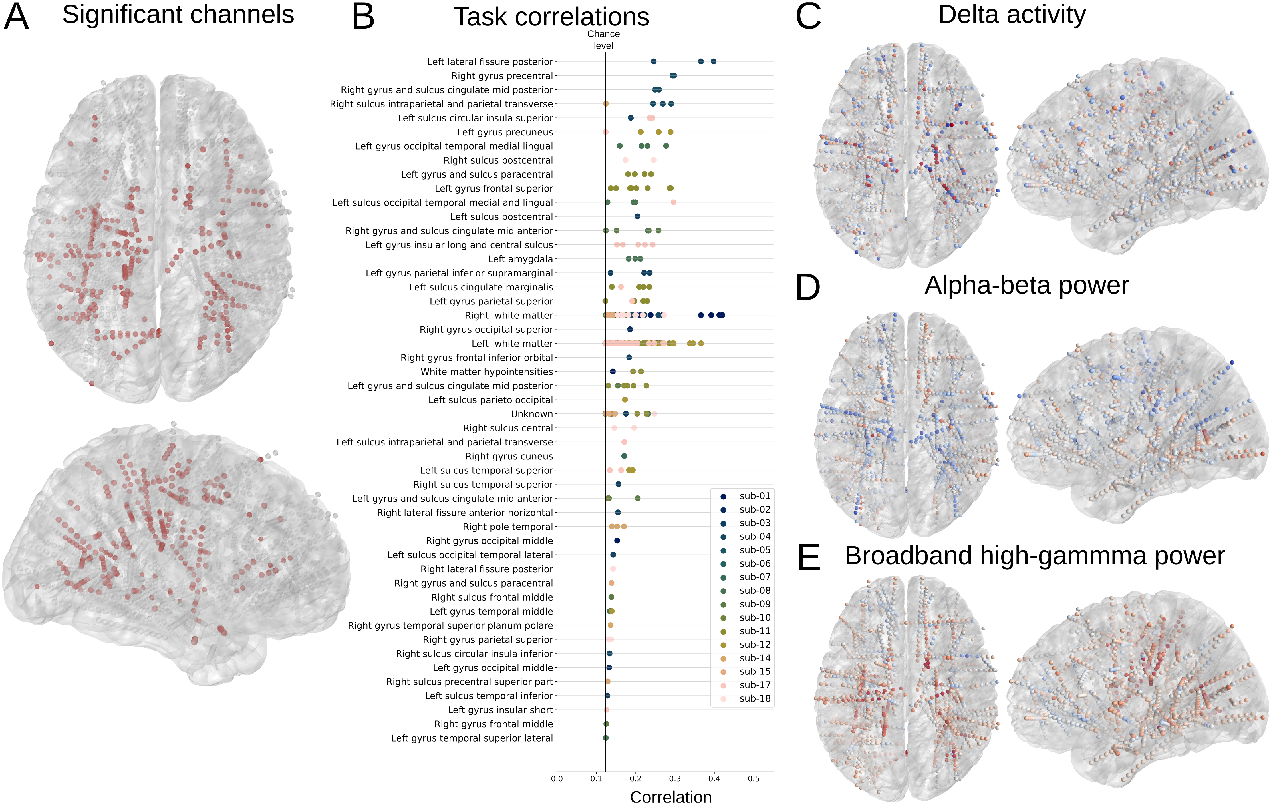
Correlations between neural activity and velocity 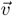 per channel. Electrode configurations of all participants are warped onto an average brain. In order to acquire the most local information, the channels are re-referenced using a Laplacian scheme and panel A and B show the correlates of broad-band high-gamma (as higher frequencies spread less far through a volume). Color scales for panel C, D and E are capped at -0.25 (blue) and 0.25 (red). **A)** Channels colored in red with a significant correlation between broadband high-gamma power and 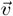. **B)** Significantly correlated channels shown in panel A separated into their anatomical locations. Sorted by average correlation per anatomical location. Anatomical locations are always determined in native space. **C)** Correlations between delta activity and velocity 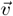. **D)** Correlations between alpha-beta power and velocity 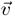. **E)** Correlations between broadband high-gamma power and velocity 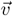. The significantly correlated channels are shown in panel A.

#### Neural data

To prepare the neural recordings, we first applied a common electrode rereference by removing the average signal of all contacts on a single electrode shaft from each contact of that shaft. In order to evaluate the spatial correlations (Figure 3), we applied a Laplacian re-reference by removing the average signal of two adjacent contacts. Next, we applied either a low, mid or high frequency filter (Figure 2). Specifically, to extract low frequencies, we applied a low-pass filter < 5 Hz (delta), for mid frequencies a band pass filter from 8 to 30 Hz (alpha & beta) and for high frequencies a band pass filter from 55 to 200 Hz (broadband high-gamma). We applied a finite impulse response filter using a hamming window, implemented in the MNE python package [40]. Finally, for mid and high frequencies, we calculated the instantaneous power using a Hilbert Transform.

#### Hand kinematics

The behavioral data were recorded with different sampling frequencies than the neural data. First, we aligned the irregularly sampled behavioral data to the neural data by finding the index of the closest matching neural timestamp for each behavioral timestamp. Next, we automatically identified gaps in the data to split them into subsets of data. No coordinates of the hand could be recorded whenever the hand was out of view, resulting in missing data. A sequence of missing data was considered a gap when four consecutive samples of behavioral data were missing. From the resulting subsets of data, we removed subsets that did not have enough data to extract five windows (as the data was windowed later). Then, for each subset, we interpolated missing behavioral samples (caused by the alignment of behavioral data to the neural data), and calculated the kinematics from the recorded coordinates, using the following terminology: the *position* is recorded in three dimensions by the motion tracker as *r*_*x*_, *r*_*y*_, *r*_*z*_. We calculated the one dimensional *distance* 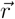 by taking the norm. As the reference frame is relative to the motion tracker, the *distance* is defined as the distance to the tracker. To calculate the *velocity v*_*x*_, *v*_*y*_, *v*_*z*_, we differentiated *r*_*x*_, *r*_*y*_, *r*_*z*_, respectively. Similarly, we calculated the non-directional *speed* 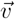 by taking the norm of the 3D velocity. To retrieve the *acceleration a*_*x*_, *a*_*y*_ and *a*_*z*_, we differentiated the velocity. To calculate the non-directional acceleration, we took the norm of the 3D acceleration. Finally, to change the reference frame to a goal-directed reference frame (Figure 4), we subtracted the cursor position from the goal position.

**Figure 4.**
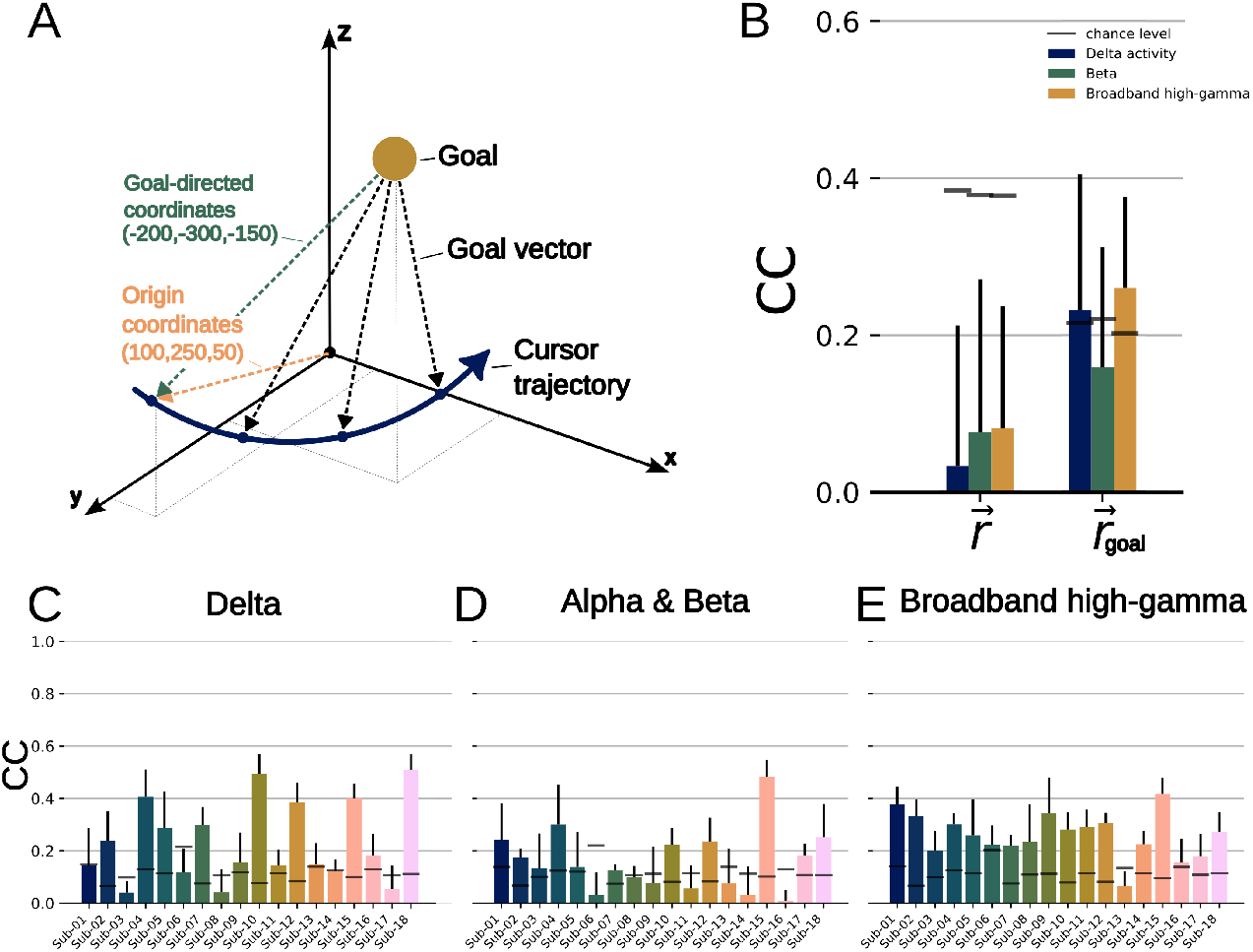
Changing the reference frame to a goal-directed reference frame allowed the decoder to decode 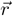 significantly above chance. **A)** To map to a goal-directed reference frame, we subtracted the coordinates of the current goal from the trajectories. By doing so, the origin of the new reference frame is located at the center of each goal. **B)** Average reconstruction correlated with 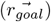 and without 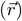 including the target vector. Decoding performance increased from below chance level to above chance level for low and high frequency information. **C)** Distance to target vector decoding performance for delta activity. **D)** Distance to target vector decoding performance for alpha-beta power. **E)** Distance to target vector decoding performance for broadband high-gamma power.

Finally, both the neural data and the behavioral data were windowed and averaged using a 300 ms window length and a 50 ms window shift and all subsets per participant were concatenated into a single dataset.

#### Feature selection

To extract the most informative features and reduce the feature space for decoding, on the training set, we selected features with the highest Pearson correlation between each channel and behavioral data. In order to ensure neural features related to all kinematics were included, we iteratively selected the best feature per kinematic until 30 unique channels were chosen.

### Decoder

To decode hand kinematics from neural data, we used the preferential subspace identification (PSID) algorithm (Figure 1d) [33]. PSID extracts behaviorally relevant dynamics that are shared between behavior and neural activity, by projecting future behavior onto past neural activity. After training, a Kalman filter predicts the movement trajectories purely based on neural activity (i.e. without ever seeing behavior in the test data; see [33] for an extensive description of PSID). PSID requires three parameters to be set: Number of latent states *N*_*x*_, number of behaviorally relevant states N_1_ and the horizon *i*. If *N*_*x*_ and *N*_1_ are different, then the remaining states (*N*_*x*_ - *N*_1_) will capture the behaviorally irrelevant dynamics, that is, neural dynamics that are not related to movements. Since we were not interested in behaviorally irrelevant dynamics, we kept *N*_*x*_ and *N*_1_ the same. Horizon *i* sets the number of future/past samples that PSID during the projection steps of the model training. A larger horizon, assuming there is enough training data to keep the projection step accurate, may lead to a better estimation at the cost of longer computation time.

First, we split the dataset for the participant using five-fold cross-validation. Then, we selected 30 channels using the selection method described in section *feature selection* on the training set. To find the optimal PSID parameters, we split the training set using a four-fold inner cross-validation, on which we performed an exhaustive grid search over *N*_1_ = [3, 5, 10, 20, 30] and *i* = [5, 10, 25, 50]. The set of parameters with the highest average performance over the inner folds was then used to train the decoder on the outer training folds. The decoding performance was evaluated by the Pearson correlation between the predicted kinematics on the test fold and the actual kinematics, called the reconstruction correlation.

### Chance level

To determine the chance level of the kinematics, we performed *n* = 1000 random circular shift permutations by splitting the actual kinematics between 10% and 90% and swapping the two sections around. The permuted kinematics should not be correlated with the neural data or with the original kinematics anymore, but still exhibit the same temporal characteristics. For each permutation, we correlated the permuted kinematics and the actual kinematics. Chance level was determined by selecting the 95th percentile of the resulting distribution of absolute correlations. To correct for multiple testing when aggregating results over participants or over channels, we took the highest chance level over the group, e.g. the highest chance level observed per kinematic over all participants.

## 3. Results

We recorded 662 ± 384 seconds (mean, std) of data per participant over 18 participants, capturing 102 ± 76 targets (mean, std), resulting in a total of 3 hours and 19 minutes of recordings. The participants completed the trials in a median of 3.3 seconds (Figure 2a). The trajectories covered the full 3D space of the hand tracker’s bounding box (Figure 2b). A stereotypical movement trajectory consisted of a large movement to the target followed by smaller corrective movements to adjust to the target position (Figure 2c). The final correction tended to be in *r*_*z*_ direction. The goals were presented sequentially within this bounding box, restricting the potential length for two sequential targets. Thus, the resulting distribution of distances between targets was skewed towards shorter distances (Figure 2d)

The electrode locations over all participants combined covered the whole brain (Figure 1a, Supplementary Figure 5). In total, 1903 contacts covered 122 unique brain areas. 1088 were located in the left hemisphere, whereas 815 were in the right hemisphere. Most contacts were labeled as left (544) or right (432) hemisphere white matter, or as unknown (128). The grey matter areas covered most were left hippocampus (n=38), left superior temporal sulcus (n=28) and superior frontal gyrus (n=23). Significant correlations were observed in 68 unique areas. See supplementary Figure 5 for a comprehensive overview.

### Hand kinematics can be decoded from brain-wide areas using low, mid and high frequency information

On average, our decoder was able to reconstruct hand movement speed 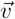 and non-directional acceleration 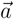 significantly above chance using delta activity, alpha-beta power and broadband high-gamma power as input (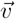 and 0.36 ± 0.16, 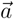 and 0.33 ± 0.15 correlation coefficient, CC, Figure 2e), with delta activity being most predictive. The mean reconstruction correlation of acceleration 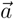 was lower than that of speed 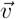, but the same performance ratio is observed between the three frequency components. The decoder was able to decode velocity in for *v*_*x*_ and *v*_*z*_ using delta activity (0.18 ± 0.14 and 0.18 ± 0.14), but not any other directional kinematic or position 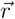. The variability of the decoding performance between participants was high, particularly when using low frequency information (Figure 2f-k). The highest decoding performance was achieved when decoding speed for *Sub-18* using delta activity (0.76 ± 0.03, Figure 2f).

### Neural correlates of hand movement are distributed throughout the brain

The neural correlates of the best performing kinematic, namely speed 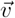, were distributed throughout the brain (Figure 3a). To extract the most local neural information, we applied a Laplacian re-reference and evaluated broadband high-gamma power neural correlates, as high frequency signals spread less far than lower frequencies. We observed significantly correlated channels throughout the brain (Figure 3a), including frontal, parietal, temporal, occipital and deeper structures. In total, broadband high-gamma power was significantly correlated with speed 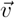 in 49 unique brain areas (Figure 3b). With the same Laplacian re-referencing scheme, low, mid and high frequency components showed correlates throughout the brain as well, (Figure 3c-e). Delta activity correlates ranged from negative (−0.48 CC) to positive (0.55 CC) correlation (Figure 3c). The mid frequency power, alpha and beta, correlates were mostly negative and ranged from -0.39 to 0.32 CC (Figure 3d), whereas the high frequency power (broadband high-gamma) showed mostly positive correlations (−0.26 to 0.42 CC, Figure 3e).

### Position could only be decoded with a goal-directed reference frame

Our decoder failed to decode the position *r*_*x*_, *r*_*y*_, *r*_*z*_ and distance 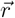 from the neural data. However, changing the original motion tracker reference frame to a goal-directed reference frame (Figure 4a) allowed the decoder to decode position and distance to the goal (Figure 4b). After changing the reference frame, the reconstruction correlation of 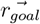 increased from below chance to significantly above chance for low (0.03 ± 0.14 to 0.23 ± 0.17 CC) and high (0.08 ± 0.16 to 0.26 ± 0.12 CC) frequency information (Figure 4b). The decoder was able to reconstruct 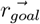 above chance: 13 of 18 participants for low frequencies, 11 of 18 participants for mid frequencies and for 17 out of 18 participants for high frequencies (Figure 4c-e).

## 4. Discussion

Movement-related neural activity has been observed throughout the human brain, but the neural contents are yet to be understood. Here, we demonstrate that goal-directed hand movements can be decoded from distributed areas, by decoding neural activity into 12 different kinematic parameters of movement. In particular, non-directional speed 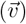 and acceleration 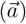 can be decoded for nearly any electrode configuration using low, mid and high frequency information. Additionally, we find weak but sufficient neural correlates to decode hand velocity (*v*_*x*_, *v*_*y*_, *v*_*z*_). Finally, we demonstrate that using a goal-directed reference frame enables the decoder to predict the position significantly above chance.

The neural correlates of low, mid and high frequency information with movement speed 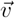 were located throughout the brain, particularly parietal, temporal, occipital and deeper brain structures such as the basal ganglia and insula. These results corroborate with the evidence that neural correlates of goal-directed behavior can be decoded from many different individual areas, as both trial-based sEEG studies [11, 12, 14, 15, 17–19, 27] and continuous motor decoding studies [16, 22, 24, 25] report successful decoding from a wide range of different frequencies and anatomical locations. In this work, frequencies across the spectrum were sufficient to decode movement, but particularly low frequency phase and high-gamma power yielded the highest decoding results. When the power of low-frequency information was used, the decoding performance dropped to chance level (not shown). The decoder in Combrisson et al. (2024) also did not select any delta power features during motor execution, supporting the importance of low-frequency phase [19] during on-going movements. Moreover, low frequency activity was among the non-power features selected most often [19]. The authors suggest a connection with the readiness potential, a slow build-up during the preparation phase, which discharges at movement initiation. Although we did not include a preparatory phase in this work, results from decoding executed movements suggest a significant relationship with on-going movement, as well [19, 26]. A similar relationship has been described in the motor cortex, initially by Schalk et al. (2007) [32], termed the local motor potential (LMP). The authors acquired the LMP by applying a 1/3s moving average signal on a raw ECoG signal, effectively applying a low-pass filter to the signal, similar to this work (< 5*Hz*) and Combrisson et al. (2024) (< 1.5*Hz*). In this work we observe that, after applying a Laplacian re-reference, the neural correlates with low frequency information remain distributed throughout the brain, suggesting that the local motor potential may be less local than it initially appeared to be. The evidence so far suggests that low frequency phase information is present throughout the brain and related to ongoing movements.

We provide a comprehensive assessment whether goal-directed behavior can continuously be decoded from brain-wide recordings, and extend previous sEEG or DBS decoding studies that reported continuous decoding results for individual movement parameters, such as force [22, 25], hand flexion [24], speed [16]. These studies mostly use trial-based paradigms with cued levels of behavior (e.g. cued levels of force). The paradigm of Shah et al. (2018) is perhaps most similar to this work, where continuous voluntary force levels were decoded with reconstruction correlations up to 0.79 cc using neural data from the subthalamic nucleus [25]. Our results show that similar results can be achieved throughout the brain, particularly when decoding speed 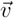 trajectories, using a naturalistic paradigm where the participant moved at their own preferred speed. The directional velocities *v*_*x*_ and *v*_*z*_ showed weaker representation in the neural data but could still be decoded above chance. Other directional kinematics could not be decoded from distributed recordings. Our decoder was able to decode *v*_*x*_ and *v*_*z*_ significantly above chance using delta activity, but on average only achieved a reconstruction correlation of 0.18 CC. Individually, the decoder performance of *v*_*x*_ and *v*_*z*_ varied from -0.01 CC to 0.45 CC. As with the other kinematics, the neural sources of directional velocities *v*_*x*_ and *v*_*z*_ are distributed throughout the brain. Neural correlates of movement direction in humans from local field potentials have been reported in the motor cortex [31]. Outside of the motor cortex, only one sEEG-based decoding study reported significant decoding of four direction 2D center-out task, with decoding accuracies ranging from 44% to 80%, using electrodes in the (pre-)motor area and the posterior supplementary motor area [19]. Expanding to ECoG, directional movement information has been decoded from non-motor areas as well, using high-gamma and the local motor potential [26]. Altogether, our results suggest that weak but sufficient directional velocity information exists outside of the motor cortex, but not for other kinematics.

Finally, both 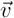 and 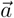 could be decoded successfully, yet 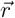 remained a challenge for the decoder. Initially, 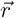 was in allocentric coordinates, meaning the reference frame was centered at the motion tracker. By transforming the coordinates into a goal-directed space (Figure 4a), the decoding performance improved from substantially below chance to above chance level (Figure 4b). On average, the three frequency bands achieved equal performance. Individually, delta achieved the highest performance, whereas the decoding performance using broad-band high-gamma power was more consistent (Figure 4c-e). Despite broadband high-gamma power being a localized signal, the decoder achieved above chance decoding for nearly all participants. Without the transformation to goal-directed space, only one out of eighteen participants achieved above chance decoding. These results suggest that movement is represented in the brain in relation to the goal. Although decoding performance improves significantly, it does require knowledge on the location of the goal. Theoretically, the selected target may be extracted directly from neural activity, but this is currently not applicable in real-world applications of brain-computer interfaces. Future work could aim to uncover new strategies to infer the user-selected goal from neural or external sources in order to apply goal-directed information to real-world applications. In spite of this, a goal-directed reference frame can already be used to rapidly calibrate neural decoder, as for example shown in Brandman et al. (2018) [41], where a unit vector pointing towards the target is used as a surrogate measure for the hidden intended cursor velocity state of a point and click system.

Using low frequency activity may increase the risk that a non-neural source related to movement causes a systematic bias in the neural data that could drive the decoding performance. However, we significantly reduced this risk by applying either a common electrode reference or a Laplacian re-reference. Any non-neural movement artifact is expected to occur in many electrodes and should thus be extracted by the re-referencing schemes. Moreover, similar decoding performance was achieved using broadband high-gamma power, where only high-frequency information is included.

Finally, the feature selection method was designed to select the features such that all twelve kinematics are represented in the selection. The feature selection method iteratively selected the top correlated feature per kinematic until a total of 30 unique features are reached. However, some channels are the top correlated for multiple kinematics. When selecting the final unique channels, some kinematics may be overrepresented due to this overlap, providing more information for that kinematics. Similarly, the decoding performance of an individual kinematic may be improved by selecting features related only to that specific kinematic.

All reported decoding results were causal decoding performances using linear decoders. Exploring non-linearly encoded behavior information and encoding of behavior in future neural data (i.e. non-causal decoding) across brain regions is an interesting future direction.

## 5. Conclusion

Here, we demonstrate that continuous goal-directed movement can be decoded from brain-wide areas using low, mid and high frequency information, and provide further evidence that the whole brain is involved during movement. Furthermore, our results suggest that movement is represented in a goal-directed context in brain-wide motor-related neural activity. These findings provide a continuous metric of ongoing movement from widespread areas in the brain, which may have implications for closed-loop brain-computer interfaces. Moreover, a robust decodable movement signal outside of the motor cortex may pave the way for the next generation of neural decoders that can be used by individuals with a compromised motor cortex, for example due to stroke.

## 6. Data and code availability

Data used in this work is openly accessible at https://osf.io/e23v9/ (accessible at publication). Code used to generate the results in this work can be found at: https://github.com/mottenhoff/decoding-continuous-goal-directed-movements

## 7. Acknowledgements

The authors would like to thank our participants for their time and effort and all involved staff at Kempenhaeghe for their invaluable support during recordings. This work is supported by the UTAP grant from Stichting de Weijerhorst and the Kootstra Talent Fellowship (2024-I). CH acknowledges funding by the Dutch Research Council (NWO) through the research project ‘Decoding Speech In SEEG (DESIS)’ with Project Number VI.Veni.194.021 and by the Kavli Foundation.

## 8. Author contributions

Conceptualization: MC, CH; Data curation: MC, MV, SG, CH; Formal analysis: MC, CH; Funding acquisition: PK, CH; Investigation: MC, MV; Methodology: MC, OS, CH; Project administration: MC, PK, CH; Resources: MV, SG, ST, HD, PK, CH; Software: MC, OS, MS; Supervision: OS, MS, PK, CH; Validation: MC, CH; Visualization: MC; Writing – original draft: MC, CH; Writing – review & editing: MC, MV, SG, ST, HD, MS, OS, PK, CH;

## 10. Supplementary material

**Figure 5.**
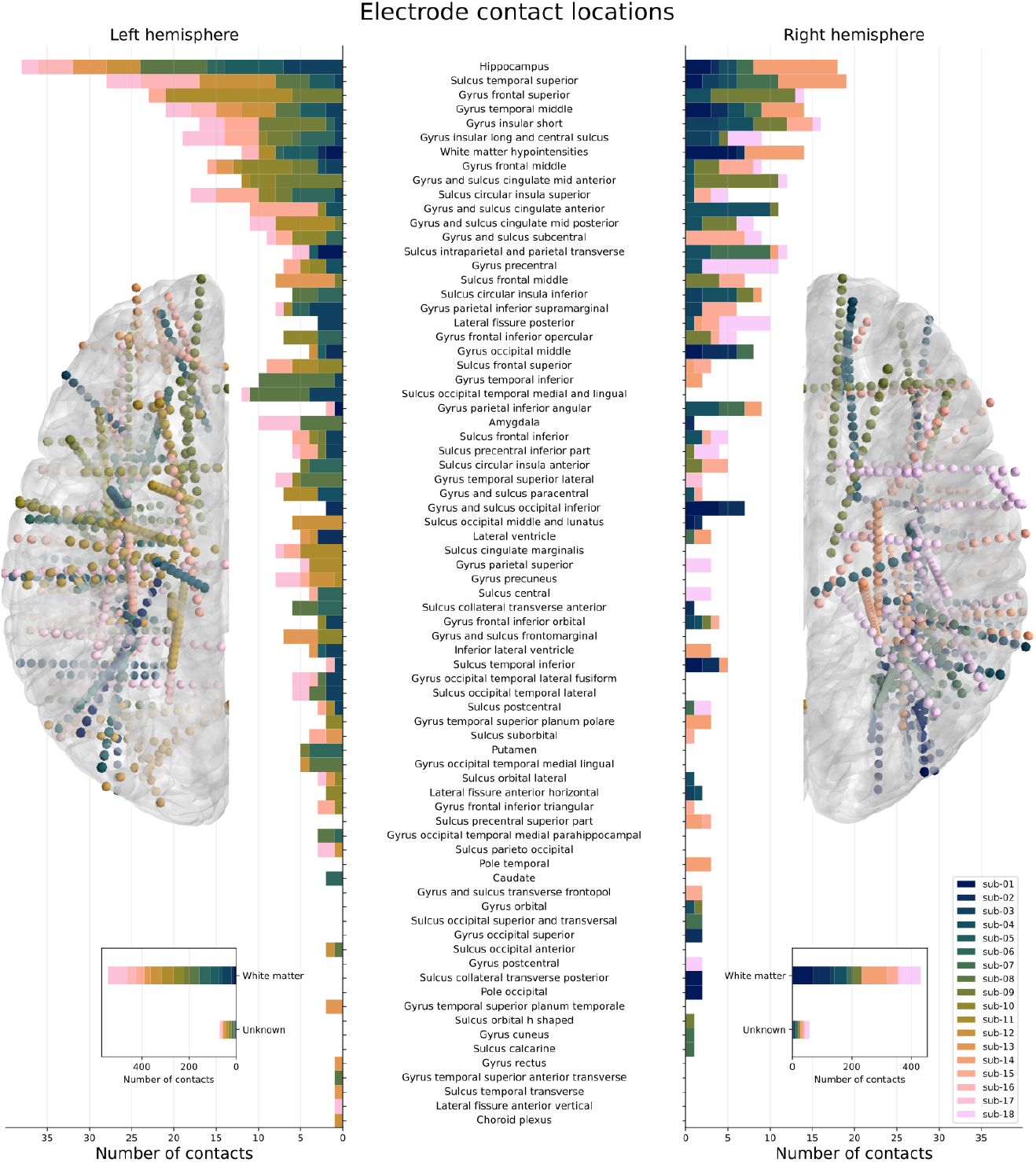
Anatomical locations covered by the electrode configurations of all participants. The insets show the contacts labeled as white matter or unknown.

## References

[1] G. J. Snoek et al. “Survey of the needs of patients with spinal cord injury: impact and priority for improvement in hand function in tetraplegics”. en. In: Spinal Cord 42.9 (Sept. 2004). Number: 9 Publisher: Nature Publishing Group, pp. 526–532. doi: 10.1038/sj.sc.3101638.

[2] Kim D. Anderson. “Targeting Recovery: Priorities of the Spinal Cord-Injured Population”. In: Journal of Neurotrauma 21.10 (Oct. 2004). Publisher: Mary Ann Liebert, Inc., publishers, pp. 1371–1383. doi: 10.1089/neu.2004.21.1371.

[3] Kim D. Anderson. “Consideration of user priorities when developing neural prosthetics”. en. In: Journal of Neural Engineering 6.5 (Sept. 2009), p. 055003. doi: 10.1088/1741-2560/6/5/055003.

[4] Leigh R. Hochberg et al. “Reach and grasp by people with tetraplegia using a neurally controlled robotic arm”. In: Nature 485.7398 (2012). Publisher: Nature Publishing Group, pp. 372–375. doi: 10.1038/nature11076.

[5] B. Wodlinger et al. “Ten-dimensional anthropomorphic arm control in a human brain-machine interface: Difficulties, solutions, and limitations”. In: Journal of Neural Engineering 12.1 (2015). Publisher: IOP Publishing. doi: 10.1088/1741-2560/12/1/016011.

[6] . International Brain Lab et al. A Brain-Wide Map of Neural Activity during Complex Behaviour. en. Pages: 2023.07.04.547681 BioRxiv. July 2023. doi: 10.1101/2023.07.04.547681.

[7] Javier Gonzalez-Castillo et al. “Whole-brain, time-locked activation with simple tasks revealed using massive averaging and model-free analysis”. In: Proceedings of the National Academy of Sciences 109.14 (Apr. 2012). Publisher: Proceedings of the National Academy of Sciences, pp. 5487–5492. doi: 10.1073/pnas.1121049109.

[8] Nicholas A. Steinmetz et al. “Distributed coding of choice, action and engagement across the mouse brain”. en. In: Nature 576.7786 (Dec. 2019). Number: 7786 Publisher: Nature Publishing Group, pp. 266–273. doi: 10.1038/s41586-019-1787-x.

[9] Carsen Stringer et al. “Spontaneous Behaviors Drive Multidimensional, Brain-wide Activity”. In: Science 364.6437 (Apr. 2019), p. 255. doi: 10.1126/science.aav7893.

[10] Misha B. Ahrens et al. “Brain-wide neuronal dynamics during motor adaptation in zebrafish”. en. In: Nature 485.7399 (May 2012). Number: 7399 Publisher: Nature Publishing Group, pp. 471–477. doi: 10.1038/nature11057.

[11] Maarten C. Ottenhoff et al. “Decoding executed and imagined grasping movements from distributed non-motor brain areas using a Riemannian decoder”. In: Frontiers in Neuroscience 17.1283491 (2023). doi: 10.3389/fnins.2023.1283491.

[12] Maarten C Ottenhoff et al. “Global motor dynamics - Invariant neural representations of motor behavior in distributed brain-wide recordings”. en. In: Journal of Neural Engineering 21.5 (Oct. 2024). Publisher: IOP Publishing, p. 056034. doi: 10.1088/1741-2552/ad851c.

[13] Richard A. Andersen, Tyson Aflalo, and Spencer Kellis. “From thought to action: The brain–machine interface in posterior parietal cortex”. In: Proceedings of the National Academy of Sciences 116.52 (Dec. 2019). Publisher: Proceedings of the National Academy of Sciences, pp. 26274–26279. doi: 10.1073/pnas.1902276116.

[14] Guangye Li et al. “Assessing differential representation of hand movements in multiple domains using stereo-electroencephalographic recordings”. en. In: NeuroImage 250 (Apr. 2022), p. 118969. doi: 10.1016/j.neuroimage.2022.118969.

[15] Meng Wang et al. “Enhancing gesture decoding performance using signals from posterior parietal cortex: a stereo-electroencephalograhy (SEEG) study”. en. In: Journal of Neural Engineering 17.4 (Sept. 2020). Publisher: IOP Publishing, p. 046043. doi: 10.1088/1741-2552/ab9987.

[16] MacAuley Smith Breault et al. “Non-motor Brain Regions in Non-dominant Hemisphere Are Influential in Decoding Movement Speed”. In: Frontiers in Neuroscience 13.JUL (July 2019), pp. 1–13. doi: 10.3389/fnins.2019.00715.

[17] Macauley Smith Breault et al. “Neural Correlates of Internal States that Capture Movement Variability”. In: 2019 41st Annual International Conference of the IEEE Engineering in Medicine and Biology Society (EMBC). ISSN: 1558-4615. July 2019, pp. 534–537. doi: 10.1109/EMBC.2019.8856778.

[18] Brian A. Murphy et al. “Contributions of subsurface cortical modulations to discrimination of executed and imagined grasp forces through stereoelectroen-cephalography”. In: PLoS ONE 11.3 (2016), pp. 1–21. doi: 10.1371/journal.pone.0150359.

[19] Etienne Combrisson et al. “Human Local Field Potentials in Motor and Non-Motor Brain Areas Encode Upcoming Movement Direction”. In: Communications Biology 7.1 (Apr. 2024), pp. 1–13. doi: 10.1038/s42003-024-06151-3.

[20] Sarah K. Wandelt et al. “Decoding grasp and speech signals from the cortical grasp circuit in a tetraplegic human”. en. In: Neuron (Mar. 2022). doi: 10.1016/j.neuron.2022.03.009.

[21] Christian Herff, Dean J. Krusienski, and Pieter Kubben. “The Potential of Stereotactic-EEG for Brain-Computer Interfaces: Current Progress and Future Directions”. English. In: Frontiers in Neuroscience 14 (2020). Publisher: Frontiers. doi: 10.3389/fnins.2020.00123.

[22] Xiaolong Wu et al. “Decoding Continuous Kinetic Information of Grasp from Stereo-electroencephalographic (SEEG) Recordings”. en. In: Journal of Neural Engineering (2022). doi: 10.1088/1741-2552/ac65b1.

[23] Syed A. Shah, Huiling Tan, and Peter Brown. “Continuous force decoding from deep brain local field potentials for Brain Computer Interfacing”. In: International IEEE/EMBS Conference on Neural Engineering, NER (2017). ISBN: 9781538619162, pp. 371–374. doi: 10.1109/NER.2017.8008367.

[24] Chad Bouton et al. “Decoding Neural Activity in Sulcal and White Matter Areas of the Brain to Accurately Predict Individual Finger Movement and Tactile Stimuli of the Human Hand”. In: Frontiers in Neuroscience 15 (2021).

[25] Syed Ahmar Shah et al. “Towards Real-Time, Continuous Decoding of Gripping Force from Deep Brain Local Field Potentials”. In: IEEE Transactions on Neural Systems and Rehabilitation Engineering 26.7 (2018). Publisher: IEEE, pp. 1460–1468. doi: 10.1109/TNSRE.2018.2837500.

[26] Aysegul Gunduz et al. “Differential roles of high gamma and local motor potentials for movement preparation and execution”. In: Brain-Computer Interfaces 3.2 (Apr. 2016). Publisher: Taylor & Francis eprint: https://doi.org/10.1080/2326263X.2016.1179087, pp. 88–102. doi: 10.1080/2326263X.2016.1179087.

[27] Alexander P. Rockhill et al. “Stereo-EEG recordings extend known distributions of canonical movement-related oscillations”. en. In: Journal of Neural Engineering 20.1 (Jan. 2023). Publisher: IOP Publishing, p. 016007. doi: 10.1088/1741-2552/acae0a.

[28] Etienne Combrisson et al. “Human local field potentials in motor and non-motor brain areas encode upcoming movement direction”. en. In: Communications Biology 7.1 (Apr. 2024). Publisher: Nature Publishing Group, pp. 1–13. doi: 10.1038/s42003-024-06151-3.

[29] Michael J. Kahana. “The Cognitive Correlates of Human Brain Oscillations”. en. In: Journal of Neuroscience 26.6 (Feb. 2006). Publisher: Society for Neuroscience Section: Mini-Review, pp. 1669–1672. doi: 10.1523/JNEUROSCI.3737-05c.2006.

[30] Lawrence M. Ward. “Synchronous neural oscillations and cognitive processes”. English. In: Trends in Cognitive Sciences 7.12 (Dec. 2003). Publisher: Elsevier, pp. 553–559. doi: 10.1016/j.tics.2003.10.012.

[31] K. Jerbi et al. “Inferring hand movement kinematics from MEG, EEG and intracranial EEG: From brain-machine interfaces to motor rehabilitation”. en. In: IRBM. NUMÉRO SPÉCIAL : LE CERVEAU DANS TOUS SES ÉTATS 32.1 (Feb. 2011), pp. 8–18. doi: 10.1016/j.irbm.2010.12.004.

[32] G. Schalk et al. “Decoding two-dimensional movement trajectories using electrocorticographic signals in humans”. en. In: Journal of Neural Engineering 4.3 (June 2007), p. 264. doi: 10.1088/1741-2560/4/3/012.

[33] Omid G. Sani et al. “Modeling behaviorally relevant neural dynamics enabled by preferential subspace identification”. en. In: Nature Neuroscience 24.1 (Jan. 2021). Number: 1 Publisher: Nature Publishing Group, pp. 140–149. doi: 10.1038/s41593-020-00733-0.

[34] C Kothe. Lab Streaming Layer (lsl). 2014.

[35] Joaquin Amigo-Vega et al. T-Rex: sTandalone Recorder of EXperiments; An easy and versatile neural recording platform. Oct. 2022. doi: 10.1101/2022.10.26.513822.

[36] Christophe Destrieux et al. “Automatic parcellation of human cortical gyri and sulci using standard anatomical nomenclature”. eng. In: NeuroImage 53.1 (Oct. 2010), pp. 1–15. doi: 10.1016/j.neuroimage.2010.06.010.

[37] Liberty S. Hamilton et al. “Semi-automated Anatomical Labeling and Inter-subject Warping of High-Density Intracranial Recording Electrodes in Electrocorticography”. English. In: Frontiers in Neuroinformatics 11 (2017). Publisher: Frontiers. doi: 10.3389/fninf.2017.00062.

[38] Fabio Crameri. Scientific colour maps. Language: eng. June 2023. doi: 10.5281/zenodo.8035877.

[39] Fabio Crameri, Grace E. Shephard, and Philip J. Heron. “The misuse of colour in science communication”. en. In: Nature Communications 11.1 (Oct. 2020). Number: 1 Publisher: Nature Publishing Group, p. 5444. doi: 10.1038/s41467-020-19160-7.

[40] Alexandre Gramfort et al. “MEG and EEG data analysis with MNE-Python”. English. In: Frontiers in Neuroscience 7 (Dec. 2013). Publisher: Frontiers. doi: 10.3389/fnins.2013.00267.

[41] David M. Brandman et al. “Rapid calibration of an intracortical brain–computer interface for people with tetraplegia”. en. In: Journal of Neural Engineering 15.2 (Jan. 2018). Publisher: IOP Publishing, p. 026007. doi: 10.1088/1741-2552/aa9ee7.

